# Hibernation site dimensions influence bat abundance and diversity in the boreal zone

**DOI:** 10.64898/2026.08.18.745443

**Authors:** Mariia Matlova, Anna Sofia Blomberg, Eeva-Maria Tidenberg, Mikhail Basok, Thomas Lilley, Mari Aas Fjelldal

## Abstract

Hibernation is a critical phase in the annual cycle of boreal bats, especially near the northern limit of their range where winter conditions are severe and suitable hibernacula are limited. In Finland, anthropogenic structures, such as abandoned bunkers, commonly serve as hibernacula for bats. In this research, we investigate how the size characteristics of bunkers affect species composition and abundance of hibernating bats. The length of the bunker was the best predictor of the presence and abundance of bats. The longest sites (∼25-26 m) had the highest probability to host all studied species, or group of species of bats, specifically *Eptesicus nilssonii*, *Plecotus auritus*, and *Myotis* bats. *Eptesicus nilssonii* appeared to be a generalist species, utilizing a variety of bunkers regardless of their length or temperature regime. Still, its number per bunker was relatively low. The highest numbers of *Myotis* bats were observed in the longest bunkers with stable temperature. The number of *P. auritus* was positively related to the bunker length; however, it was rare in the studied hibernacula. Different species showed different patterns in hibernacula space usage. *Eptesicus nilssonii* hibernated in any part of the bunker, while *Myotis* bats used the most remote from the entrances areas in small sites, and occupied the whole area of the longest bunkers.

## Introduction

In the boreal zone, bats face various challenges associated with annual fluctuations in temperature, which are greater than in any other ecozone. Short and moderately warm summers are followed by long, cold winters with the seasonal temperature range being as great as 50°C. Besides low temperatures, winter is characterized by frozen waterbodies and prolonged snow cover, which dramatically change the environment and force animals to develop a range of overwintering strategies depending on their ecology and physiology. For instance, considering that most of the insects overwinter as eggs or in a dormant phase (Bale and Hayward, 2010), insectivorous animals, such as bats, lose their food source when environmental conditions decline in the autumn (Speakman and Rowland, 1999). Some bat species are capable of long-distance migration and thus avoid the long winters (Fleming, 2019). However, resident bats depend on hibernation as a strategy to survive the unfavourable winter conditions (Geiser, 2013).

Hibernation is characterized by prolonged bouts of torpor during which the metabolic rate and body temperature are low, as well as changes in other physiological processes resulting in minimized energy expenditure (Geiser, 2004; Dzal and Brigham, 2013). The main advantage of hibernation is a decrease in energy and water expenditure during a time when it is difficult to replenish these resources (Boyles et al., 2019). Furthermore, bats congregate at particular overwintering sites, where mating also occurs, thus contributing to the relevance of these sites as ‘genetic hotspots’ (Fraser and McGuire, 2023).

However, hibernating bats are vulnerable to various threats. First and foremost, they risk starvation as insect abundance is reduced to near zero at northerly latitudes (Perry, 2013); therefore, the overwinter survival of bats relies on fat reserves accumulated before winter (Beer and Richards, 1956; Reusch et al., 2023; Fjelldal et al., 2024). Most of these reserves are spent on short-term arousals throughout the hibernation to offset accumulated physiological costs caused by metabolic depression during deep, prolonged torpor (Thomas et al., 1990; Humphries et al., 2003; Landes et al., 2020). Torpid bats are also less capable of detecting and responding to predator presence due to their reduced metabolism. Aggregations of inactive bats have become a food source for predators, e.g. mustelids (Reimer and Willis, 2025), red foxes, raccoons (Cichocki et al., 2021), as well as for owls (Sommer et al., 2009), wood mice (Haarsma and Kaal, 2016), brown rats (Gloza-Rausch et al., 2025) and even great tits (Estók et al., 2010). In some cases, hibernating bats comprise up to 96% of a predator’s diet (Cichocki et al., 2021).

Winter temperatures below zero centigrade increase the likelihood of freezing (Gillette and Kimbrough, 1970) and inability to replenish water reserves, forcing bats to find appropriate overwintering shelters. Decades of research indicate that temperature conditions are of vital importance for successful hibernation in bats e.g. (Gaisler, 1970; Raesly and Gates, 1987; Webb et al., 1996; Perry, 2013; De Bruyn et al., 2021). There is no ‘optimal’ hibernation temperatures; bats utilize a wide range of temperatures during their hibernation period (Masing and Lutsar, 2007; Boyles et al., 2017). Yet, there are important restrictions, as the surrounding temperatures determine torpor bout length and therefore strongly affect the energetics of hibernating bats (Boyles et al., 2019; Villada-Cadavid et al., 2025). Both too cold and too hot microclimates can lead to increased energy expenditure in hibernating bats (Hope and Jones, 2012). More stable microclimate in hibernacula are associated with greater energy savings than less stable (Newman et al., 2024).

Besides low temperatures, winters bring an inability to replenish water reserves, as waterbodies become covered with ice and snow. Bats are particularly susceptible to dehydration because their wing membranes provide a large surface area for water evaporation (Thomas and Cloutier, 1992). To avoid death from dehydration while hibernating, bats tend to use hibernacula with high humidity. As it has been shown in experiments, water depletion in hibernating bats can be compensated only if relative humidity exceeds 99.3% (Thomas and Cloutier, 1992). Some studies suggest that humidity may play a more important role in overwintering success than other abiotic factors, as arousal frequency, and hence, energy consumption, is inversely related to humidity (Ben-Hamo et al., 2013). Species characterized by small body weight can die of dehydration sooner than of starvation (Speakman and Racey, 1989). Despite the obvious benefits of high humidity for bats at hibernation sites, high humidity may also promote the growth of pathogenic microbiota, including causative agents of deadly bat diseases (Marroquin et al., 2017). Therefore, choosing hibernacula with appropriate microclimatic conditions is always a trade-off.

Temperature and humidity preferences vary both between species and within the same species from one individual to another. Among boreal bat species, some tend to hibernate at relatively high temperatures and humidity, for example the European species *Myotis dasycneme* (forearm 43-49 mm, weight 13-18 g), *M. mystacinus* (forearm 32-36.5 mm, weight 4-7 g), and the North-American species *Myotis sodalis* (forearm 34-43 mm, weight 5-11 g), *M. septentrionalis* (forearm 34-38 mm, weight 5-8 g) (Webb et al., 1996; Caceres and Barclay, 2000; Masing and Lutsar, 2007; Dietz and Kiefer, 2016; Brack et al., 2025; Roby et al., 2025). On the other end of the spectrum, there are cold-hardy species that are observed at relatively low temperatures, such as *Eptesicus nilssonii* (forearm 37.1-44.2 mm, weight 9-13 g), *E. fuscus* (forearm 39-54 mm, weight 11-23 g) and *Plecotus auritus* (forearm 35.5-42.8 mm, weight 6-9 g) (Kurta and Baker, 1990; Webb et al., 1996; Masing and Lutsar, 2007; Siivonen and Wermundsen, 2008; Dietz and Kiefer, 2016). Moreover, bats actively select the most suitable microclimate by changing their position within the hibernacula during arousals (Daan, 1973; Masing, 1987). However, an understanding of species-specific optimal hibernation conditions within local ecoregions is still lacking (Boyles, 2017; Ryan et al., 2019). Thus, the question of what affects the preferences of bats when selecting hibernation sites is of current interest.

The variety of hibernacula used by bats is great; however, in temperate climates the chosen hibernation sites should be well protected against both opportunistic predators and freezing winter temperatures. Various natural and anthropogenic undergrounds, such as caves (Zukal et al., 2017), mines (Rydell et al., 2018), tunnels (Beer and Richards, 1956), abandoned military buildings (Kokurewicz et al., 2019), cellars (Vintulis and Pētersons, 2014), as well as deep rock crevices (De Boer et al., 2013), rock outcrops and boulder fields (Blomberg et al., 2025), serve as suitable overwintering shelters. With a substantial body of research manifests hibernation sites to be of a great importance (Mitchell-Jones et al., 2007; Voigt et al., 2014; Van Der Meij et al., 2015), we still know very little on how bats select overwintering shelters, and which characteristics affect their choice.

In Finland, situated in Northeast Europe, the abandoned bunkers built along the eastern border in 1940s are known as bat hibernation sites with three of them listed as the most important bat hibernacula in the country (Eurobats, 2016). Although there are more than 700 military constructions in the defense line (Beck et al., 2003), only a number of them located in the southern part of the country has been examined for the presence of bats. Examination showed a notable divergency in bat abundance: from 1 or 2 bats in some of the bunkers to several dozen in the most abundant ones (Wermundsen and Siivonen, 2010a). Five species of bats commonly hibernate there: *Eptesicus nilssonii*, *Myotis brandtii*, *M. mystacinus*, *M. daubentonii* and *Plecotus auritus*. Additional to this, there are three species that may also potentially hibernate there, however, they are rarely observed: *M. nattereri* occurs yearly at a handful of locations in southwestern Finland, and for *M. dasycneme* and *E. serotinus*, only individual record exist (Tidenberg et al., 2019). Recent studies have shown that *Pipistrellus nathusii*, a species considered as a long-distant migrant, also hibernate in Finland, a geographic area more associated with summer breeding grounds for the species (Blomberg et al., 2025, 2021). However, we still know very little about hibernation ecology of these species in Finland.

The overwhelming majority of studies on the topic are focused on temperature and humidity conditions (Webb et al., 1996; Brack et al., 2025). Only some research include dimensional characteristics of a hibernaculum, such as volume, length, number of available rooms as well as the number of hiding possibilities showing they can affect the abundance of bat species (Kim and Yoo, 2004; De Boer et al., 2013). In this work we investigate the effects of dimensional characteristics of the hibernacula on bat species composition and abundance in boreal zone. Our work was based on the following two hypotheses: i) the number of species and their abundance will increase with the size of the hibernacula as larger hibernacula will offer bats a wider range of microclimates, given that the notion of “size’’ is properly defined; ii) species will be split into two groups: generalized species that utilize any type of the hibernacula regardless of its dimensional characteristics, and other species demanding specific characteristics of hibernacula.

## Materials and methods

### Study site

The study was conducted in Virolahti and Miehikkälä municipalities, located in South-Eastern Finland (Figure 1). The hibernacula examined included bunkers and anthropogenic caves that are part of the extensive Salpalinja fortification line, constructed between 1940 and 1944 along the Finnish-Russian border. We examined 69 hibernacula, out of which nine are anthropogenic caves made in granite rock, and the rest are built of concrete. The majority of them are one-floor buildings, some have vertical turrets that form the second and sometimes the third floors. For convenience, all sites are regarded as bunkers in this study.

**Figure 1.**
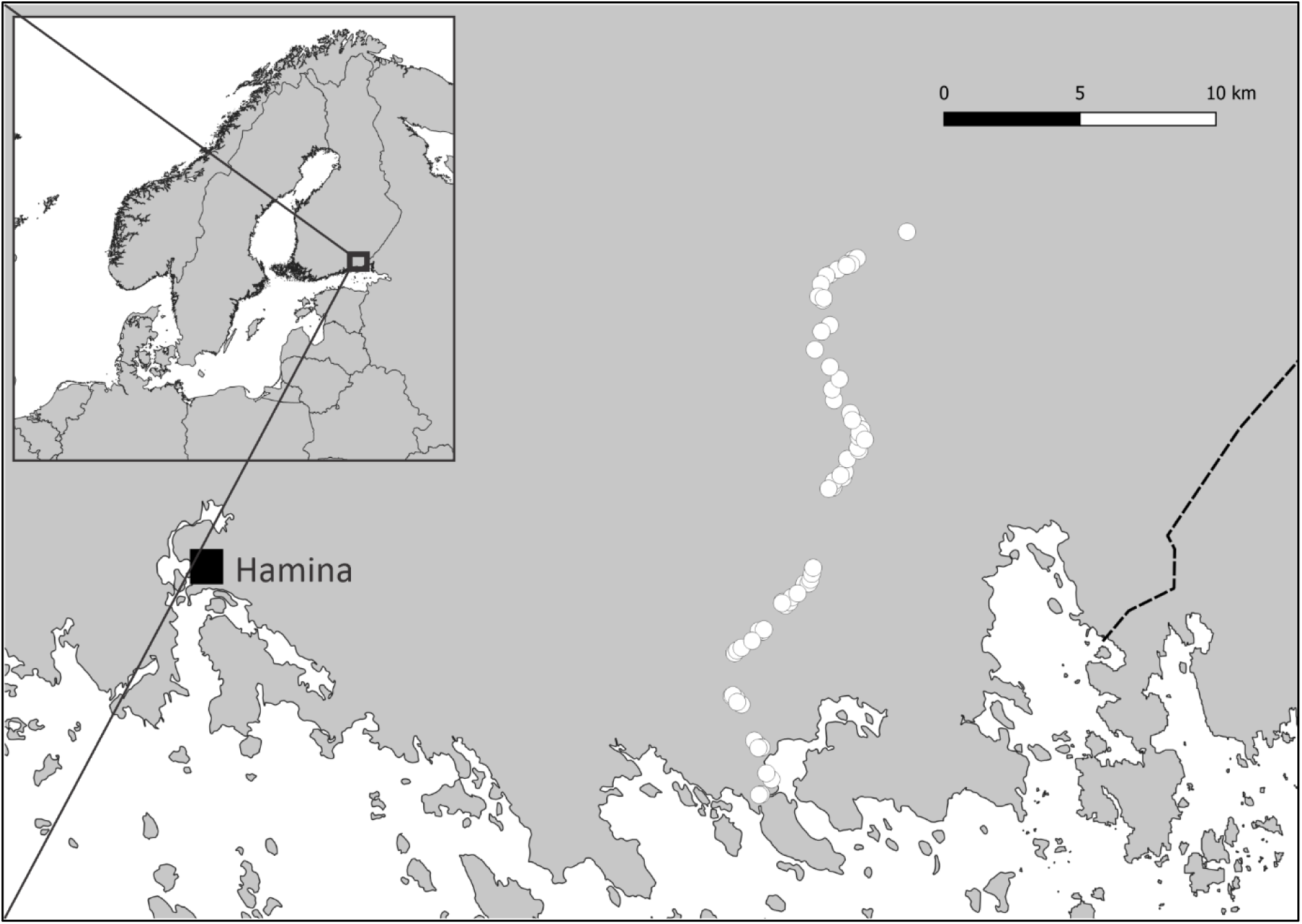
Map of study area. White dots represent bunker locations.

### Dimensional features of the bunkers

To get dimensional characteristics of study sites, we measured each site using a laser distance measurer tool (Cocraft HD 400-2, working distances 0.2 – 40 m, accuracy ±1.5 mm). While in the field, a schematic hand-written layouts of the study sites with all measurements were created. Those layouts were digitized later. In total, we derived measurements from 69 bunkers. Further calculations of the area, perimeter, volume, and bunker length were done with an assistance of a computer program written in Python (the code uploaded to online Zenodo-repository, DOI: https://doi.org/10.5281/zenodo.21984710).

The average height of concrete bunkers was ∼ 2 m, the height of the cave-like bunkers varied from 2 to 5 m. The area of the smallest and the largest bunkers were 15 m^2^ and 300 m^2^ respectively (mean 57.3 m^2^) with cave-like constructions being the largest ones. The volume ranged from 26 m^3^ to 1143 m3; the perimeter ranged from 26 m to 221 m. The length was calculated as the distance between the most remote location inside the bunker and the nearest entrance. The minimum length of the bunker was 5.5 m, the maximum length was 26 m. We regarded all openings, including doorways, shooting windows and ventilation holes, as entrances. Usually, a bunker had one or two doorways and one or two shooting windows. Thus, the maximum number of entrances per bunker in our study system was four.

### Bat census

Monitoring of hibernating bats at these sites has been ongoing since 2007. In this study, we used data derived during 2013-2025. Each February, the Finnish Museum of Natural History organizes the monitoring effort, which is primarily carried out by a dedicated group of volunteers. The number of surveyed sites has grown since the early years and now typically ranges between 80 and 100, depending on snow conditions and site accessibility. We focused on 69 bunkers that were most often accessible and were visited during most of the censuses. During each census, data were collected on the number of bats, their species, and the specific locations of individuals or clusters. In 2022 – 2025, data on bat distribution within the surveyed bunkers were obtained as well.

Species identification was based on morphological characteristics and excluded any bat handling. In this study, *M. daubentonii*, *M. brandtii* and *M. mystacinus* were combined into *Myotis* group. Two latter species are often combined into one group, *M. brandtii/ mystacinus*, due to difficulties with separating them from each other during winter censuses when no bat handling can be applied. In our case, we had to put *M. daubentonii* into the joint *Myotis* group as well because over the bat census years there are accumulated number of cases when bats were classified to genus *Myotis* only.

### Temperature in the bunkers

In winter 2024 – 2025, we put temperature loggers (DS1921G-F5 Thermochron ibuttons, accuracy ± 1°C) into fifteen bunkers that differed in their size characteristics. The loggers were located in the innermost parts of the bunkers, at the height of 1.5-2.5 m. Due to some issues (some ibuttons disappeared from the sites) our sample included data from fourteen bunkers collected during January and February 2025.

### Statistical analysis

All analyses were performed in R (version 4.4.1). To understand how the different dimensional parameters of a bunker impact bat communities (see response variables tested in the sections below), we were interested in the following predictors: area of the bunker (m^2^); perimeter (m); volume (m^3^); the number of entrances; the length of the bunker (m).

Prior to model fitting, we tested for correlation between the explanatory variables. The area was heavily positively correlated with the volume (cor: 0.98) and perimeter (cor: 0.94). To avoid multicollinearity, highly correlated predictors were not included in the same model. We fitted generalized linear mixed models using lme4 package (Bates et al., 2015). Initially, both location and year were included as random effects. However, the variance associated with year was estimated to be negligible in all models. Therefore, year was excluded from the final models, and bunker ID was retained as the only random effect. Model selection was based on Akaike information criterion (AIC), where the model with the lowest AICc value was considered the best fit (Aho et al., 2014). We used ggplot2 and marginaleffects packages (Arel-Bundock et al., 2024) to visualize the results of the models.

### The effect of dimensional characteristics of hibernacula on probability of bat occurrence

For each bunker and census year, the presence or absence of each bat species was calculated. Species presence was coded as 1 when at least one individual of a given species was detected in a bunker, and as 0 otherwise. To investigate the effect of dimensional characteristics on probability of encountering bats of different species, we fit five binomial generalized linear mixed-effects models with dimensional features as predictors, presence-absence of different species as the response variable, and bunker ID as a random effect (see Table S1 in Supplementary Materials for model list).

### The effect of dimensional characteristics of hibernacula on bat abundance

To determine the effect of dimensional characteristics of the hibernaculum on the abundance of bats of different species, we performed five negative binomial generalized linear mixed-effects models with the dimensional characteristics as predictors, the number of bats of different species as the response variable, and bunker ID as a random effect (Table S2 in Supplementary Materials).

### The effect of mean temperature and temperature stability on bat abundance

To investigate the effect of mean temperature in the innermost part of the bunkers on the abundance of bats of different species, we used data on temperature collected during January and February 2025, and data on bat abundance collected during February 2025. Temperature stability was calculated as the difference between maximum and minimum temperature recorded in each bunker. We fitted two negative binomial generalized linear mixed-effects models. The first one included mean temperature and bunker length as predictors, number of bats of different species as response variable, and bunker ID as a random effect. The second one included temperature stability as predictor, number of bats of different species as response variable, and bunker ID as a random effect. The number of individuals of *P. auritus* registered in 2025 was very low (11 individuals), thus this species was excluded from the final models.

### The effect of dimensional characteristics of bunkers on distribution of bats

The distance from each bat to the nearest entrance was calculated with assistance of the computer program written in Python. To compare the spatial distribution of bats within bunkers of different sizes, we calculated a relative distance as the ratio of the distance from the entrance at which an individual was recorded to the bunker length. To investigate the effect of dimensional characteristics of the hibernaculum on the bat distribution, we performed five binomial generalized linear mixed-effects models with the dimensional characteristics as predictors, the relative distance of bats of different species as the response variable, and bunker ID as a random effect (Table S3 in Supplementary Materials). The number of individuals of *P. auritus* was very low, thus this species was excluded from the final models.

## Results

### Bat counts and size characteristics of the bunkers

Out of the 69 measured bunkers, between 36 and 69 bunkers were visited annually during the 13 winter censuses (Table 1). The annual total number of bats ranged from 187 to 327, consisting of three species or species groups: *Myotis* bats (this group included *M. daubentonii*, *M. brandtii* and *M. mystacinus*) and *E. nilssonii* were the most abundant species, while *P. auritus* was much less numerous.

**Table 1.** Overview of the number of visited bunkers, and the number of bats of different species counted during winter censuses 2013-2025.

| Year | Number of visited bunkers | Total number of bats | Number of bats of different species |  |  |  |
| --- | --- | --- | --- | --- | --- | --- |
|  |  |  | *Eptesicus nilssonii* | *Myotis sp.* | *Plecotus auritus* | Not identified |
| 2025 | 69 | 323 | 151 | 158 | 11 | 3 |
| 2024 | 65 | 239 | 99 | 128 | 9 | 3 |
| 2023 | 59 | 283 | 117 | 156 | 8 | 2 |
| 2022 | 61 | 281 | 102 | 169 | 7 | 3 |
| 2021 | 61 | 289 | 105 | 173 | 11 | 0 |
| 2020 | 63 | 327 | 125 | 190 | 12 | 0 |
| 2019 | 53 | 240 | 77 | 153 | 10 | 0 |
| 2018 | 48 | 246 | 86 | 148 | 9 | 3 |
| 2017 | 36 | 187 | 64 | 110 | 9 | 4 |
| 2016 | 49 | 216 | 83 | 128 | 5 | 0 |
| 2015 | 50 | 208 | 79 | 121 | 7 | 1 |
| 2014 | 47 | 265 | 92 | 168 | 5 | 0 |
| 2013 | 47 | 232 | 66 | 160 | 6 | 0 |

### Predicted probability of observing bats of different species

The model that best explained the effect of the size of a hibernaculum on the probability of observing a bat included the length of the bunker as a predictor in interaction with bat species, with bunker ID as a random effect. The predicted probability of observing *E. nilssonii* was high across all bunker lengths; even in the bunker with the smallest length (5.5 m), the predicted observation probability exceeded 50% and increased significantly (p < 0.001) to almost 100% with increasing length of the hibernacula (Fig. 2; Table 2a). The predicted probability of observing *Myotis* bats and *Plecotus auritus* also increased with increasing bunker length; however, the observed patterns differed significantly from the one shown for *E. nilssonii* (p < 0.001). In the bunkers whose length was less than 10 m, the predicted probability of observing a *Myotis* bat was low (less than 10%). When the length exceeded 15 m, the probability began to increase steeply and reached almost 100% in the longest bunkers (> 25 m). The pattern for *P. auritus* resembled the one shown by *Myotis* bats. However, even in the longest bunkers the predicted probability of observing *P. auritus* was merely ∼50%, and the confidence intervals were large (Fig. 2).

**Figure 2.**
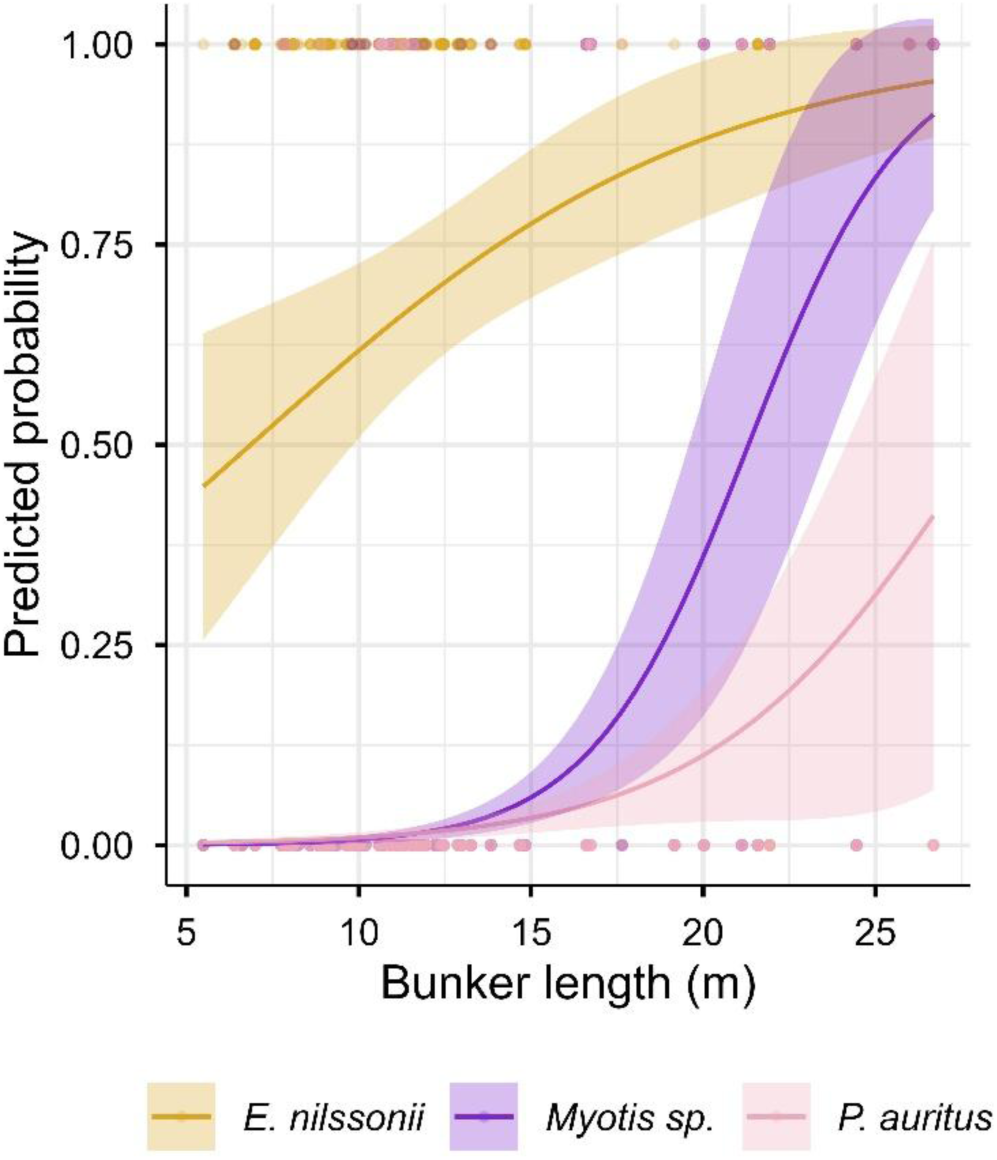
The predicted effects (lines and confidence intervals) of bunker length on the probabilities of observing bats of different species (group of species): *E. nilssonii*; *Myotis* bats; *P. auritus*. The datapoints indicate actual observations.

**Table 2.** Model results presenting the final best-fit model explaining: (a) effect of bunker length on probability of bat occurrence; (b) effect of bunker length on bat abundance; (c) effect of mean temperature on bat abundance; (d) effect of temperature stability on bat abundance; (e) effect of bunker length on bat distribution within the bunkers.

| Variable | Estimate (SE) | p-value | z-value |
| --- | --- | --- | --- |
| a) Effect of bunker length on probability of bat occurrence |  |  |  |
| Bunker ID (random effect) | 2.28 (SD = 1.51) |  |  |
| Intercept (*E. nilssonii*) | -0.6 (0.69) | 0.3686 | -0.899 |
| Intercept (*Myotis sp.*) | -8.8 (0.8) | <0.001 | -10.8 |
| Intercept (*P. auritus*) | -6.3 (0.75) | <0.001 | -8.4 |
| Length x *E. nilssonii* | 0.15 (0.05) | 0.005 | 2.9 |
| Length x *Myotis sp.* | 0.33 (0.06) | <0.001 | 5.96 |
| Length x *P. auritus* | 0.1 (0.05) | 0.0237 | 2.3 |
| b) Effect of bunker length on bat abundance |  |  |  |
| Bunker ID (random effect) | 0.76 (SD = 0.87) |  |  |
| Intercept (*E. nilssonii*) | -0.31 (0.33) | 0.357 | - 0.921 |
| Intercept (*Myotis sp.*) | -6.96 (0.32) | <0.001 | -22.145 |
| Intercept (*P. auritus*) | -4.96 (0.37) | <0.001 | -13.474 |
| Length x *E. nilssonii* | 0.05 (0.03) | 0.06 | 1.875 |
| Length x *Myotis sp.* | 0.36 (0.02) | <0.001 | 21.513 |
| Length x *P. auritus* | 0.14 (0.02) | <0.001 | 7.092 |
| c) Effect of mean temperature and bunker length on bat abundance |  |  |  |
| Bunker ID (random effect) | 0.03 (SD = 0.17) |  |  |
| Intercept (*E. nilssonii*) | 0.57 (0.498) | 0.256 | 1.136 |
| Intercept (*Myotis sp.*) | -7.17 (1.27) | <0.001 | -5.66 |
| Length x *E. nilssonii* | 0.02 (0.03) | 0.482 | 0.703 |
| Length x *Myotis sp.* | 0.03 (0.05) | 0.582 | 0.551 |
| Mean temperature x *E. nilssonii* | 0.03 (0.08) | 0.712 | 0.37 |
| Mean temperature x *Myotis sp.* | 1.84 (0.37) | <0.001 | 5.04 |
| d) Effect of temperature stability on bat abundance |  |  |  |
| Bunker ID (random effect) | 0.12 (SD = 0.34) |  |  |
| Intercept (*E. nilssonii*) | 0.97 (0.33) | <0.001 | 2.93 |
| Intercept (*Myotis sp.*) | 5.22 (0.6) | <0.001 | 8.767 |
| Temperature range x *E. nilssonii* | -0.01 (0.07) | 0.855 | -0.183 |
| Temperature range x *Myotis sp.* | -2.23 (0.37) | <0.001 | -6.05 |
| e) Effect of bunker length on bat distribution within the bunkers |  |  |  |
| Bunker ID (random effect) | 0 (SD = 0) |  |  |
| Intercept (*E. nilssonii*) | 1.14 (0.27) | <0.001 | 4.242 |
| Intercept (*Myotis sp.*) | 2.36 (0.76) | <0.001 | 3.087 |
| Length x *E. nilssonii* | -0.05 (0.02) | <0.001 | -2.76 |
| Length x *Myotis sp.* | -0.05 (0.04) | 0.143 | -1.464 |

### The relationship between bunker length and bat abundance

The best model to explain the relationship between dimensional characteristics of a hibernaculum and the abundance of bats of different species occupying them included the length of the bunker as a predictor, the number of bats of different species as a response variable, and bunker ID as a random effect. The length of the bunker positively influenced the number of hibernating bats of all three species (or group of species in case of *Myotis* bats), although the effect size differed across species (Fig. 3; Table 2b). The number of *E. nilssonii* hibernating simultaneously in the same bunker ranged from zero to ten (mean number of individuals per bunker: 1.76 ± 1.79 SD), and the predicted number of individuals increased only slightly with the length of the bunker (Table 2b; Fig. 3).

**Figure 3.**
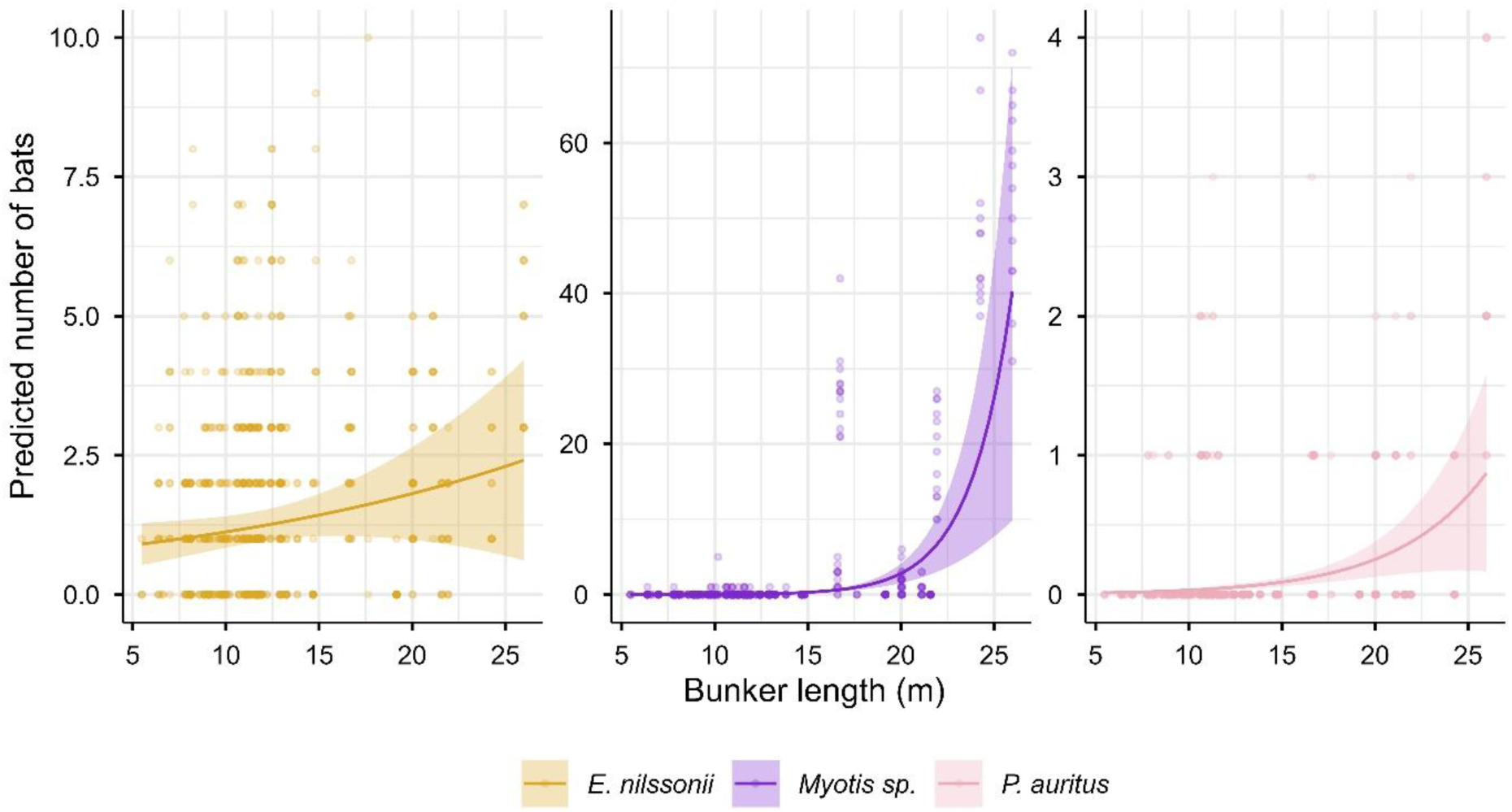
The predicted effect (lines and confidence intervals) of bunker length on the abundance of different bat species (group of species): *E. nilssonii*; *Myotis* bats; *P. auritus*. The datapoints indicate actual observations.

The number of *Myotis* bats hibernating in the same hibernacula ranged from zero to 74 (mean: 2.77 ± 10.5 SD), and was strongly affected by the length of the bunker (Table 2b; Fig. 3). In the small and middle-size bunkers which length did not exceed 15 m, one could occasionally observe one *Myotis* bat. Only once in 13 years of monitoring did we find five individuals hibernating in one of the small bunkers. Meanwhile, if the length of the hibernacula exceeded ∼15 m, the number of *Myotis* bats increased noticeably with their abundance varying from 20 up to 74 individuals in different bunkers. However, there were some exceptions: several bunkers with the length of 17 – 22 m never hosted a *Myotis* bat.

The general abundance of *Plecotus auritus* hibernating in the bunkers was low in comparison to the other species, with observed numbers of bats ranging from zero to 4 (mean: 0.2 ± 0.5 SD) and only a small positive effect of increasing. It is interesting to note that in one of the longest bunkers, despite the low abundance in general, this species was observed to hibernate every year without exception during the past 13 censuses. *P. auritus* was never found in the bunkers if their length was less than 7 m.

### The relation between the temperature and bat abundance

The model to explain the relations between the mean temperature and number of observed bats of different species in a smaller dataset, included the length of the bunker in addition to the mean temperature in the innermost part of the bunker as predictors, the number of bats of different species as a response variable, and bunker ID as a random effect. Mean temperatures had a positive influence on the number of Myotis bats (Fig. 4A; Table 2c). The highest abundance was observed in the bunkers where the mean temperature exceeded +4.5°C. Interestingly, the two most abundant bunkers with more than 50 and 70 bats respectively had equal temperature +5.1°C ± 0.4 SD in the innermost parts. We never observed *Myotis* bats if the mean temperature in the bunker was below zero. Additionally, we tested the effect of temperature stability in the innermost part on bat abundance. According to the model, temperature stability was a significant predictor of the abundance of *Myotis* bats (Fig. 4B; Table 2d). Most of the observed individuals were found in those bunkers where the difference between the minimum and maximum temperatures was less than 1.5°C.

**Figure 4.**
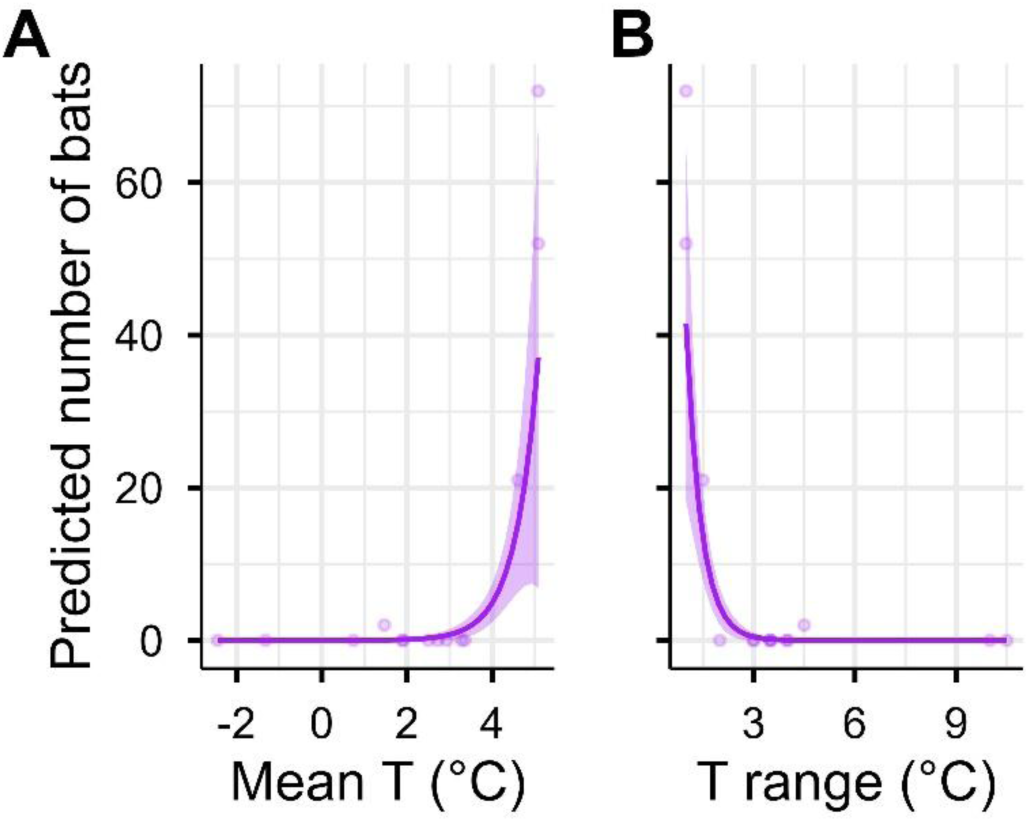
The predicted effects of the mean temperature (A) and temperature stability (B) on the number of *Myotis* bats

There was no significant effect of the mean temperature or temperature fluctuations on the number of hibernating *E. nilssonii* (Table 2c, 2d). Most of individuals were found in the bunkers with the mean temperature varying from +0.7°C ± 0.9 SD to +5.1°C ± 0.4 SD; some individuals were found in the bunker at – 2.4°C ± 2.4 SD. *E. nilssonii* usually utilized those bunkers with the temperature range of maximum 5°C. Five individuals were found in the bunker where the difference between the minimum and maximum temperatures was 10°C.

### Effect of the bunker length on bat distribution

The best model to explain the relationship between dimensional characteristics of a hibernaculum and the distribution of bats of different species occupying them included the length of the bunker as a predictor, the number of bats of different species as a response variable, and bunker ID as a random effect. Bunker length had a strong effect on how *Myotis* bats were distributed along the bunker length (Fig. 5; Table 2e). While in small sites individuals were observed in the innermost parts only, with the increase of the bunker length *Myotis* bats began to use the whole area of the hibernacula. In the longest bunkers, several individuals were hibernating in the vicinity of the entrance doorway. Solitary animals were observed in small bunkers. Small groups of *Myotis* bats were observed if the bunker length exceeded 15 m. The biggest clusters comprising of up to 20 individuals were observed in the longest bunkers.

**Figure 5.**
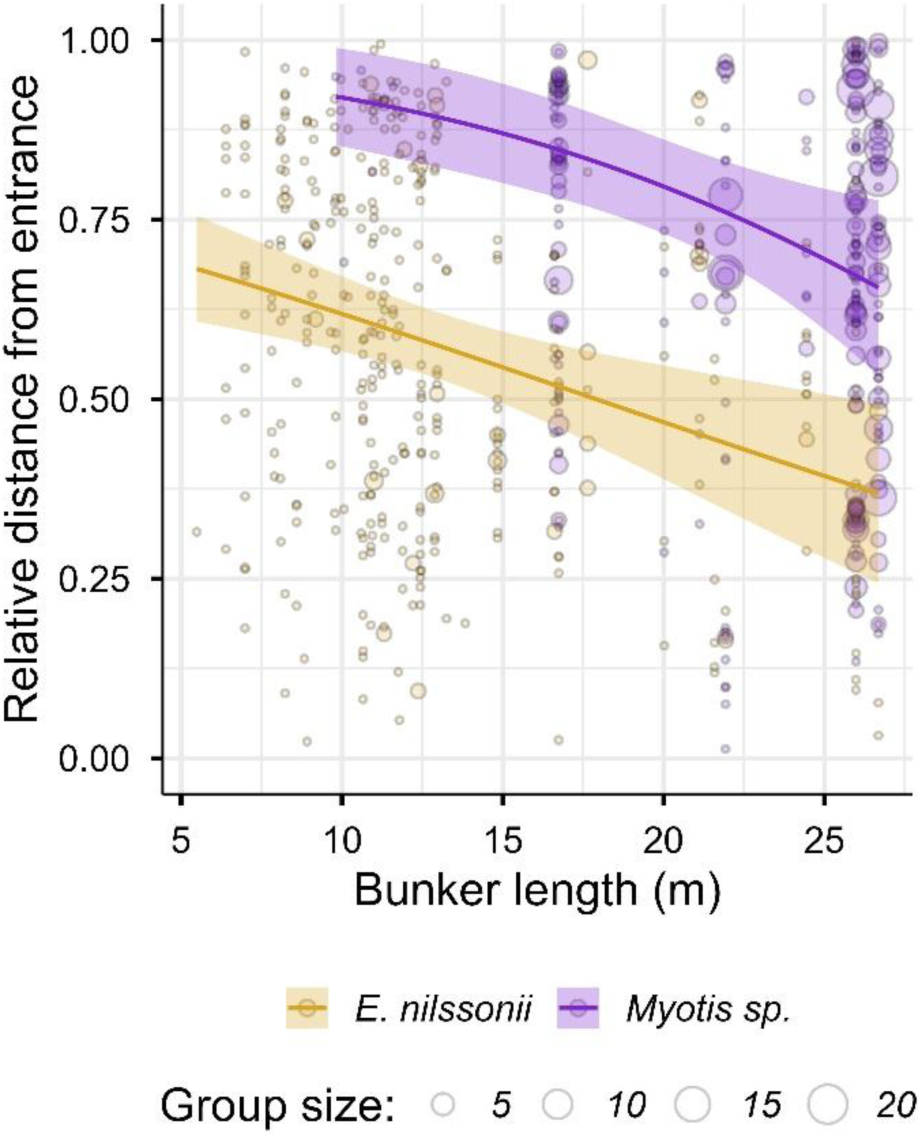
The relation between the length of the bunker and the relative distance from the entrance at which *E. nilssonii* and *Myotis* bats were observed.

The effect of the bunker length on the distribution of *E. nilssonii* was moderate (Fig. 5; Table 2e). Individuals of this species were observed at the entrance, middle, or innermost parts of the bunkers regardless of their length. However, in the longest bunkers *E. nilssonii* avoided the most remote parts and were usually observed in the middle and entrance areas. Although *E. nilssonii* never huddled into big clusters, a groups of two or three individuals were registered in all types of the bunkers.

## Discussion

This long-term study on bats hibernating in bunkers in South-East Finland describes impacts of hibernacula characteristics on bat species composition and abundance. Our results demonstrate that the length of the hibernacula (i.e., the distance from the entrance to the innermost part of the bunker) is the best predictor of presence and abundance of different species of bats in this system. The highest probability of observing all three species/group of species of bats, specifically *E. nilssonii*, *P. auritus*, and *Myotis* bats, was in the longest bunkers (∼25-26 m). The abundance of *Myotis* bats was positively affected by the length of the hibernacula, the mean temperature and the temperature stability. In contrast, there was no significant effect of the bunker-length or temperature on the number of *E. nilssonii*. The number of *P. auritus* was positively related to the length of the bunker; however, the overall abundance of this species observed in the studied bunkers was low. In small-sized bunkers, both *E. nilssonii* and *Myotis* bats tended to hibernate in the most remote area from the entrances. With the increase in the bunker length, *E. nilssonii* were more often registered in the middle parts or close to an entrance. In long bunkers (>15 m), *Myotis* bats used the whole area of the hibernacula with some individuals being observed near the entrances. The largest clusters of *Myotis* bats (∼20 individuals) were found in the longest bunkers, while clusters of *E. nilssonii* did not exceed three individuals and were observed in bunkers independent of their size. Our results contribute to filling the knowledge gap in our understanding of overwintering site selection by boreal bats facing long, harsh winters in areas where natural hibernacula (e.g., caves) are scarce.

Larger hibernacula host more bats overall, as larger sites can offer more available space and higher variety of microclimatic conditions. However, the concept of size can have multiple interpretations. Our study shows that the bunker-length is a key feature that shapes species composition and abundance of bats in our system. This measurement may reflect the gradual changes in microclimate beginning from the entrance where temperature is usually highly unstable (Daan and Wichers, 1968) and humidity low (Klüg-Baerwald et al., 2024), to the innermost point which is more buffered from the prevailing ambient conditions outside (Daan and Wichers, 1968; Baranauskas, 2003). In contrast to our expectations, we found no clear linear relation between the length and the mean temperature measured in the bunkers. The longest sites were not necessarily the warmest; however, the few bunkers in which site length was positively associated with mean temperature exhibited the highest bat abundance and species richness. The unexpected lack of correlation between bunker length and temperature could be a result of airflow patterns in bunkers with certain shapes or multiple number of entrances, which could cool or dry the hibernacula despite the size. Unfortunately, due to the limited number of bunkers, we could not investigate other dimensional features (e.g., bunker-shape or number of entrances) that may affect site selection by bats (Lesiński, 1986; Kłys, 2013). Still, our study provides a first report on species-specific hibernacula characteristic important for the occurrence and abundance of hibernating bats in Finland.

*Eptesicus nilssonii* had the highest overall occurrence probability across bunkers in our study system. The predicted probability of observing *E. nilssonii* was higher than 50% even in the smallest bunkers (∼5 m length) and increased to almost 100% in the longest bunkers. Still, the number of *E. nilssonii* per bunker was relatively low (maximum of 10 individuals in the same hibernacula) with low evidence for a positive relation between the abundance and the length of the bunker. Across its geographical species distribution range, whenever abandoned bunkers or other military buildings serve as bat hibernacula, e.g., in northwestern Russia (Chistyakov, 2009; Belkin et al., 2019), Latvia (Vintulis and Pētersons, 2014), Lithuania (Baranauskas, 2006a; Masing et al., 2009), Belarus (Godlevska et al., 2023), as well as in previous studies in Finland (Wermundsen and Siivonen, 2010a), the number of *E. nilssonii* observed per hibernaculum is relatively low regardless of hibernacula size, and comparable to what we found in our study. Even when huge underground bunker systems are available, such as the 32-km long Nietoperek bat reserve in Poland, *E. nilssonii* avoid these and instead overwinter in smaller bunkers nearby (Grzywiński et al., 2012).

The temperature conditions within the hibernacula did not appear to be the primary factor driving the overwintering site selection in our studied *E. nilssonii*; the mean temperature had no significant effect on the number of bats, and individuals were recorded across a wide range of temperature conditions. Our observations are similar to reports made on the cold-hardy North-American bat species *Eptesicus fuscus* (Klüg-Baerwald et al., 2024). According to previous studies, *E. nilssonii* and *E. fuscus* demonstrate a great tolerance towards microclimatic instability and high flexibility in microclimate selection, hibernating at temperatures ranging from −10°C to +10°C (Webb et al., 1996; Rydell et al., 2018; Belkin et al., 2021). Some studies report that *E. nilssonii* most often hibernates at temperatures between 0°C to +5°C (Strelkov, 1958; Siivonen and Wermundsen, 2008). This range of favourable temperature conditions was also observed in the bunkers studied here, which likely explains the frequent occurrence of this species in our study. Being well adapted to harsh conditions, *E. nilssonii* may even benefit from them by reducing competition, occupying short, cold-exposed bunkers that are unsuitable for thermophilic species.

The manner in which *E. nilssonii* utilises the space within our studied bunkers further reflects these adaptations. Individuals hibernated in the innermost part of short, small bunkers; in medium-sized bunkers, they occupied the entire space; while in long bunkers, individuals tended to hibernate in the front parts (with some exceptions). Thermal conditions near the entrance (e.g., small sites or entrance areas of any site) can be strongly correlated with outside temperatures (Daan, 1973; Boyles et al., 2017) thus constituting a challenging environment that only cold-hardy species, such as *E. nilssonii*, can tolerate. Hibernating close to the entrance can be beneficial as bats can use weather cues to detect foraging opportunities. Thus, *E. nilssonii* can be opportunistically active during boreal winter (Blomberg et al., 2025) and use suitable days for hunting and replenishing water. Although research on hibernacula space usage by *E. nilssonii* is scarce, a similar pattern to that observed in our study was reported in fortification systems in northwest Russia (Chistyakov, 2009). In non-military hibernacula, such as mines and pits, *E. nilssonii* uses the space according to thermal conditions: in sites with large temperature differences between entrance and inner parts, with relatively high temperature in inner sections (>6°C), individuals use near-entrance areas; in contrast, in hibernacula with relatively cool temperatures (0°C – 5°C) it can hibernate throughout the whole site regardless of hibernacula size (Strelkov, 1958; Bolʹshakov et al., 2005; Smirnov et al., 2008; Matlova et al., 2024). It is noteworthy to mention that we here discuss mid-winter (February) census-data; the patterns could therefore differ from observations made on *E. nilssonii* early or late in the winter.

For *Myotis* bats, we detected a positive relationship for both occurrence and abundance with bunker length. A similar effect of hibernacula length has been reported for the Asian species *Myotis formosus*, hibernating in abandoned gold mines (Kim and Yoo, 2004). In Europe, in general, larger hibernacula host larger populations of *Myotis* bats, with hundreds and thousands of bats hibernating in the most extensive sites, such as several kilometers long natural caves (Bolʹshakov et al., 2005; Piksa et al., 2013), artificial caves (Lutsar et al., 2000; Smirnov et al., 2007), and military undergrounds (Cichocki et al., 2015).

The selection of larger/longer hibernacula could be explained by thermal preferences of *Myotis* bats. Unlike *E. nilssonii*, *Myotis* bats across Europe (Baranauskas, 2003; Piksa et al., 2013) and North-America (Webb et al., 1996; Brack et al., 2025) tend to avoid low and frequently fluctuating temperatures with our results mirroring these findings. Still, thermal preferences of *M. daubentonii*, *M. brandtii* and *M. mystacinus* are known to differ. *Myotis daubentonii* is believed to have the widest range of selected hibernation temperatures (Masing and Lutsar, 2007; Kovalyov, 2017; Malyavina et al., 2025), with some studies describing it as a thermophilic species (Dzięciołowski et al., 2022) and others as a cold-tolerant bat (Spitzenberger et al., 2024). It is frequently reported hibernating in small, relatively cool hibernacula, such as cellars, basements, and fortifications, in contrast to *M. brandtii* and *M. mystacinus* (Lesiński et al., 2004; Ciechanowski et al., 2006; Grzywiński et al., 2012; Godlevska et al., 2023). Although relatively few studies have investigated *M. brandtii* and *M. mystacinus* separately, those that have indicate that *M. brandtii* is the most thermally stenobiontic of the two (Smirnov et al., 2008; Piksa et al., 2013). *Myotis brandtii* prefers relatively high and stable temperature (Piksa et al., 2013) and seldom appears in small hibernacula, such as little cellars (Masing, 1983; Lesiński et al., 2004; Baranauskas, 2006b) or small bunkers (Chistyakov, 2009; Grzywiński et al., 2012; Lesiński and Stolarz, 2024). This conclusion is supported by (Baranauskas, 2006a) where *M. brandtii* appeared in hibernacula after the entrance was partially blocked and the temperature and humidity increased. In contrast, *M. mystacinus* is described as an eurythermal species tolerant of low temperatures and unstable conditions, thus showing a greater range of distribution across different hibernacula (Smirnov et al., 2008; Piksa et al., 2013).

Interestingly, in areas where several relatively similar hibernacula occur in close proximity, such as karst systems or clusters of abandoned mines, species dominance varies between neighbouring sites. Hibernacula with high numbers of *M. daubentonii* generally have low numbers of *M. brandtii* and *M. mystacinus*, and vice versa; sites dominated by *M. mystacinus* also tend to contain few *M. brandtii*. This picture has been observed in the natural caves of Sudetic Mountains (Furmankiewicz and Furmankiewicz, 2002), as well as in limestone pits and sand mines (Smirnov et al., 2007; Kovalyov, 2017). A somewhat similar pattern was observed in our study area: previous research showed that *M. brandtii/mystacinus* prevailed over *M. daubentonii* in the same bunker system as presented in our study (Wermundsen and Siivonen, 2010a). However, such segregation could potentially be explained by unmeasured environmental factors. For instance, humidity plays a vital role in overwintering success of hibernating bats, affecting torpor-arousal patterns and general welfare of animals (Ben-Hamo et al., 2013; McGuire et al., 2021), and different species may prefer different humidity levels. Another explanation could be hidden in social organization of abovementioned species. Highly gregarious species that tend to hibernate in clusters, like *M. brandtii/mystacinus* (Smirnov and Vekhnik, 2009), accumulate in particular hibernacula with conspecifics. In contrast, *M. daubentonii* usually hibernates solitary or in smaller groups (Wermundsen and Siivonen, 2010b), thus being more flexible and less dependent on conspecifics when choosing suitable hibernation sites.

The distribution of *Myotis* bats within the hibernacula in our study also reflects their thermal preferences, although grouping three Myotis species with varying preferences likely obscure species-specific microclimate relationships, which should be kept in mind for the interpretation of our results. In small sites, where the microclimate often follows the changes in ambient temperature (Boyles, 2017), individuals tended to hibernate in the innermost parts, while in long bunkers, *Myotis* bats were found throughout the length of the bunker. This latter observation might be explained by the fact that we had three different species, each requiring different conditions, and hence occupying different parts of the bunkers (Smirnov et al., 2008; Matlova et al., 2024). However, it is noteworthy to consider that space distribution within hibernacula may also be a result of active site selection by individuals depending on their internal states regardless of species (Boyles et al., 2007).

*Plecotus auritus* was rare in our study system. Although the probability of encountering this species increased with the bunker length, its abundance remained low even in the longest bunkers. *Plecotus auritus* often shares hibernacula with *E. nilssonii* (Masing, 1983; Piksa et al., 2013; Vintulis and Pētersons, 2014; Rydell et al., 2018), which occupied most of the studied bunkers; thus, its low abundance was unexpected. The low abundance could be explained by the ecology of *P. auritus*, which is a sedentary species that tends to hibernate close to its summer territories (Furmankiewicz, 2016). If the area is environmentally unsuitable as a summer habitat, we might expect low numbers or even the absence of *P. auritus* in the nearby hibernacula. This interpretation is supported by studies indicating that many small sites distributed across large territories, such as root cellars, basements, or natural karst cavities can serve *P. auritus* better than a single large hibernacula (Furmankiewicz and Furmankiewicz, 2002; Lesiński et al., 2004; Grzywiński et al., 2012; Vintulis and Pētersons, 2014). Another explanation of the low abundance observed in our study area may be related to microclimate preferences of *P. auritus*. Although it exhibits traits similar to *E. nilssonii*, including the use of small hibernacula (Lesiński et al., 2004; Vintulis and Pētersons, 2014), occupancy of near-entrance areas in large sites (Smirnov et al., 2008), and tolerance of relatively cool temperatures (Webb et al., 1996; Wermundsen and Siivonen, 2010b; Malyavina et al., 2025), some researchers emphasize that *P. auritus* avoids unstable, rapidly changing microclimates and air-drafty locations (Strelkov, 1958; Nagel and Nagel, 1991; Masing and Lutsar, 2007; Piksa et al., 2013). Given that larger hibernacula often provide a more stable environment (Boyles, 2017), we might expect a higher abundance of *P. auritus* in such sites, which is consistent with the predictions of our model. Indeed, in some areas larger hibernacula host greater numbers of *P. auritus* (Masing, 1983; Lesiński and Stolarz, 2024), with the largest populations observed in large hibernacula with stable environment, such as the underground fortification system of Nietoperek bat reserve in Poland (460-630 individuals) (Urbanczyk, 1990; Cichocki et al., 2015), large artificial caves in northwestern Russia (>300 individuals) (Chistyakov, 2001), and large limestone pits in Samara region of Russia (> 1900 individuals) (Smirnov et al., 2007). Nevertheless, it is clear from our study that we still lack information on the selection of hibernacula by *P. auritus* in Finland.

Many important anthropogenic hibernacula in Europe, including abandoned bunkers, fortresses, tunnels, and mines, date back to the 19^th^ century. Decades of deterioration and weathering contribute to the degradation of these hibernacula. Under such circumstances, the conservation of existing important overwintering sites is essential. An increasing number of sites undergo renovation, which often involves the closure of existing entrances or the construction of new entrances to replace collapsed ones (Baranauskas, 2006a). However, reconstruction can lead to dramatical changes of microclimate which in turn causes changes in bat species composition. It is therefore becoming a common practice to assess key habitat features that are important predictors of bat use before interference in order to mitigate negative impacts (Johnson and Kuchta, 2024). Here, we present results showing that in boreal zone, the length of the hibernaculum can be an important predictor of bat species richness, and also predict the abundance of *Myotis* bats. We expect that our findings will help to provide recommendations for effective site management in the interest of bat conservation. Nevertheless, given that not all hibernacula of suitable length were occupied by bats, further research is needed to investigate the effects of other factors. These may include abiotic factors, such as entrance size, airflow patterns, temperature regimes throughout the site, and humidity, as well as biotic factors, including human disturbance, predation pressure, and competition between bats.

## Acknowledgements

We would like to thank Valeriia Bohodist, Asko Ijäs, Petri Asikainen, Anna-Kitty Ekstam, Juha Forsten, Niclas and Siri Fritzén, Arvo Gran, Helena Haakana, Nina Hagner-Wahlsten, Rasmus Karlsson, Antti Karppi, Sanna Karttunen, Noora Kauppila, Juho Kosunen, Pasi Kyheröinen, Kristian Lindqvist, Risto Lindstedt, Melissa Meierhofer, Kalle Meller, Katarina Meramo, Timo and Mahla Metsänen, Jacqueline Nelms, Laura Niinimäki, Andreas Otterbeck, Tiina Saalasti, Silva Sallamaa, Saku Salonen, Ella Sippola, Kati Suominen, Miina Suutari, Tanya Troitsky, Hanna Tuominen, Ville Vasko, Riitta Vikberg, Teemu Virtanen and Ralf Wahlsten for extensive help with bunker measurements and annual bat winter census. We are grateful to two anonymous reviewers for helping us greatly improve the manuscript.

## Supplementary materials

**Table S1.** The list of ranked models with relationship between dimensional characteristics of bunkers and probability of bat occurrence.

| Rank | Model | df | AIC | ΔAIC |
| --- | --- | --- | --- | --- |
| 1 | Probability ~ Species*Length + (1\| Bunker ID) | 7 | 1322.15 | 0 |
| 2 | Probability ~ Species*Perimeter + (1\|Bunker ID) | 7 | 1347.93 | 25.8 |
| 3 | Probability ~ Species*Number_of_entrances + (1\|Bunker ID) | 7 | 1354.88 | 32.7 |
| 4 | Probability ~ Species*Area + (1\|Bunker ID) | 7 | 1355.9 | 33.8 |
| 5 | Probability ~ Species*Volume + (1\|Bunker ID) | 7 | 1363.64 | 41.5 |

**Table S2.** The list of ranked models with relationship between dimensional characteristics of bunkers and bat abundance.

| Rank | Model | df | AIC | ΔAIC |
| --- | --- | --- | --- | --- |
| 1 | Bat_abundance ~ Species*Length + (1\| Bunker ID) | 8 | 3882.17 | 0 |
| 2 | Bat_abundance ~ Species*Number_of_entrances +<br>(1\|Bunker ID) | 8 | 4123.29 | 241.2 |
| 3 | Bat_abundance ~ Species*Perimeter + (1\|Bunker ID) | 8 | 4140.25 | 258.1 |
| 4 | Bat_abundance ~ Species*Area + (1\|Bunker ID) | 8 | 4205.3 | 323.1 |
| 5 | Bat_abundance ~ Species*Volume + (1\|Bunker ID) | 8 | 8895.88 | 5013.7 |

**Table S3.**
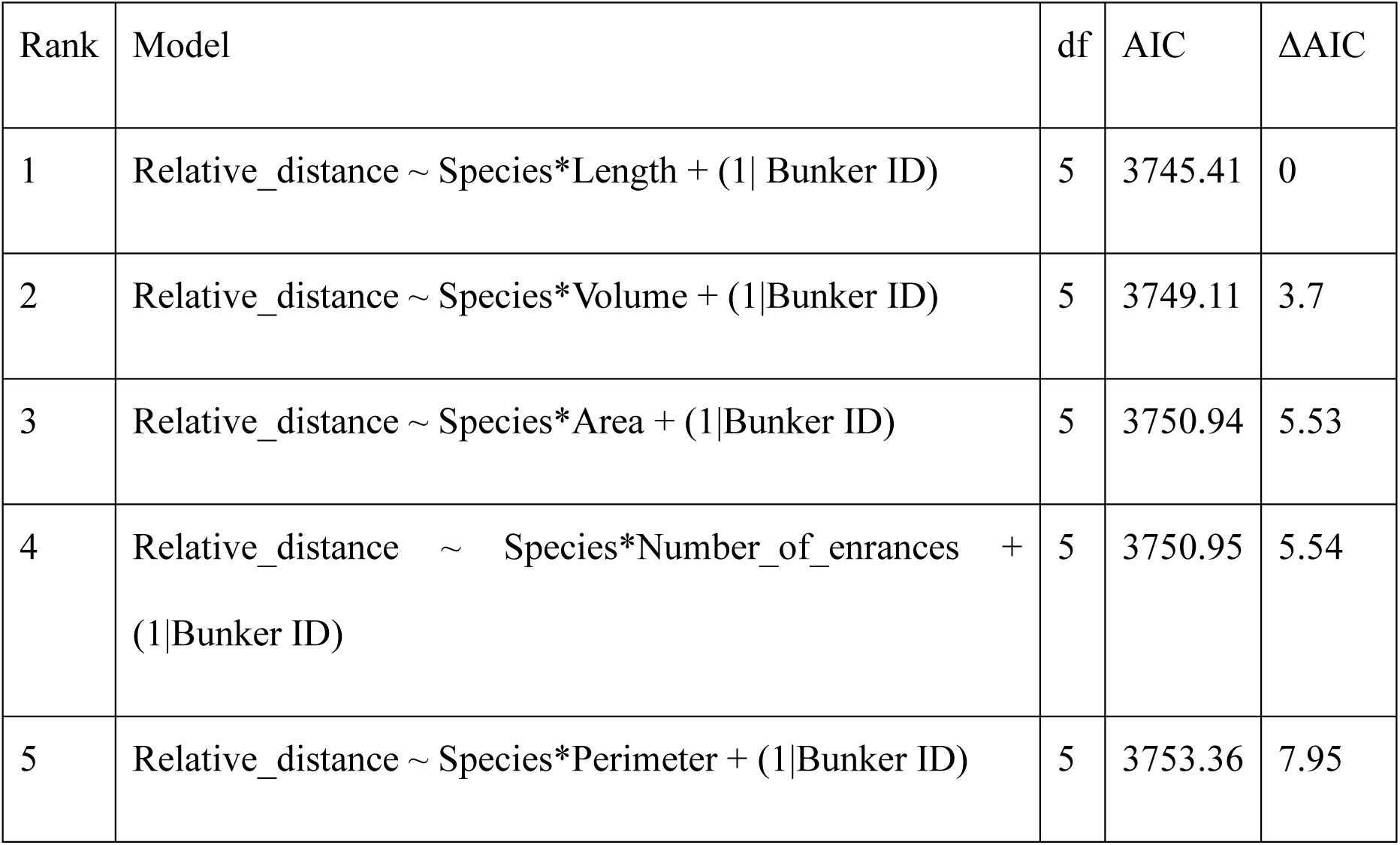
The list of ranked models with relationship between dimensional characteristics of bunkers and relative distance from the entrance at which bats were hibernating.

| Rank | Model | df | AIC | ΔAIC |
| --- | --- | --- | --- | --- |
| 1 | Relative_distance ~ Species*Length + (1\| Bunker ID) | 5 | 3745.41 | 0 |
| 2 | Relative_distance ~ Species*Volume + (1\|Bunker ID) | 5 | 3749.11 | 3.7 |
| 3 | Relative_distance ~ Species*Area + (1\|Bunker ID) | 5 | 3750.94 | 5.53 |
| 4 | Relative_distance ~ Species*Number_of_enrances +<br>(1\|Bunker ID) | 5 | 3750.95 | 5.54 |
| 5 | Relative_distance ~ Species*Perimeter + (1\|Bunker ID) | 5 | 3753.36 | 7.95 |

